# Effects of oral cannabidiol (CBD) on spontaneous opioid withdrawal in male and female rats

**DOI:** 10.1101/2025.10.08.680984

**Authors:** Bryan W. Jenkins, Cerina Pang, Robbie Y. Kuang, Elise M. Weerts, Catherine F. Moore

**Author notes:** Corresponding author: Catherine F. Moore, Ph.D. Department of Psychiatry and Behavioral Sciences, Johns Hopkins Bayview Medical Campus, Behavioral Biology Research Center 5510 Nathan Shock Drive, Suite 3000, Baltimore, MD, 21224, USA.

## Abstract

Opioid use disorder (OUD) remains a public health crisis in the United States. A key factor in continued use, relapse risk, and overdose is the severe withdrawal syndrome that accompanies abstinence. Observational studies suggest cannabis may improve outcomes for patients with OUD. Cannabidiol (CBD), a non-intoxicating compound found in cannabis, is being investigated as a potential treatment for OUD. This study investigated whether CBD alleviated withdrawal symptoms in a rat model of opioid dependence. Sprague Dawley rats (N = 100, 50% female) were administered escalating doses of morphine across 10 days (10–50 mg/kg, s.c., twice daily). Following abrupt discontinuation, withdrawal outcomes were evaluated across acute (38-hr) and protracted (up to day 7) timepoints. Rats were treated with pure CBD (10 or 30 mg/kg, p.o.) or vehicle (sesame oil; 1 mg/ml) daily, beginning 14-hrs after their final morphine or saline injection (n = 8-9 per sex/group). Withdrawal severity was assessed through physical measurements of body weight, food intake, and somatic signs (e.g., body shakes, diarrhea), and pain sensitivity, as well as measurements of anxiety-like behaviors in the protracted phase. Compared to non-dependent controls, morphine-dependent rats had decreased body weight and food intake, showed greater somatic signs, and had increased pain sensitivity that peaked in acute withdrawal (38-hr). Oral CBD did not affect physical symptoms of opioid withdrawal nor protracted anxiety-like behaviors. These data indicate that CBD alone may have limited effectiveness for treating opioid withdrawal. Reports of improved withdrawal symptoms after cannabis use may be attributed to other compounds in cannabis.

**Public health significance:** This study suggests that daily oral treatment with pure CBD at doses available to consumers does not improve physical or psychological symptoms of withdrawal from chronic opioids.

## Introduction

Approximately 120 people die every day from an opioid-related drug overdose in the United States (Mattson et al., 2021). The U.S. opioid epidemic continues to be a critical public health crisis that demands novel interventions to reduce or prevent overdose risk in patients with opioid use disorder (OUD). A key factor in continued or relapsed use, which increases overdose risk, is the emergence of a severe withdrawal syndrome following discontinuation of chronic use. The opioid withdrawal syndrome includes physical/somatic symptoms such as tremors, gastrointestinal distress, diarrhea, restlessness, muscle aches, and increased pain sensitivity (hyperalgesia) (Gipson, Dunn, Bull, Ulangkaya, & Hossain, 2021; Higgins, Smith, & Matthews, 2019; Janiri et al., 2005; Jasinski, 1981; Martin & Jasinski, 1969). Physical symptoms typically emerge within 8–12 hours of discontinued use, peak between 24–72 hours and gradually subside over 1–2 weeks (i.e., the acute phase). The opioid withdrawal syndrome also features psychological symptoms including craving, anxiety, negative affect, and anhedonia, which can persist for weeks or months beyond the acute somatic symptoms (i.e., the protracted phase) (Gipson, et al., 2021; Higgins, et al., 2019; Janiri, et al., 2005; Jasinski, 1981; Martin & Jasinski, 1969). The intense discomfort brought on by withdrawal can motivate users to alleviate symptoms by re-initiating use, which increases risk for overdose through decreased tolerance after a drug-free period (Williams, Samples, Crystal, & Olfson, 2020). Medical interventions targeting withdrawal in patients with OUD are critical for sustained cessation and decreased overdose risk.

Standard of care for managing withdrawal is medication-assisted treatment, which includes supervised cessation of nonmedical opioid use concomitant with opioid replacement therapy using an opioid agonist with reduced abuse liability (e.g., methadone or buprenorphine). However, patients who are seeking or engaged in treatment are often interested in non-opioid options, including using cannabis or cannabis-derived products for clinical management of OUD (Kudrich, Chen, Meng, Bachi, & Hurd, 2024). Cannabidiol (CBD), a non-intoxicating cannabinoid found in the *Cannabis sativa* plant, is of particular interest for OUD medications development (Hurd et al., 2015; Kudrich, et al., 2024). CBD exerts bioactive effects through interactions via multiple molecular mechanisms of action. This includes cannabinoid type 1 (CB1R) and type 2 (CB2R) receptors and opioid (mu, delta) receptors, serotonin 5HT1A, transient receptor potential cation channel, TRPV1 ion channels and GABA_­_receptors as well as enzymes that control endocannabinoid synthesis, reuptake and degradation (e.g., fatty acid amide hydrolase, FAAH; peroxisome proliferator-activated receptors, PPARs). (Bakas et al., 2017; de Almeida & Devi, 2020; Kathmann, Flau, Redmer, Trankle, & Schlicker, 2006; McPartland, Duncan, Di Marzo, & Pertwee, 2015; Ruffolo et al., 2022; Russo & Marcu, 2017). The broad receptor engagement of CBD underlies its proposed wide-ranging uses as a therapeutic, including as an anxiolytic, analgesic, anti-depressant, anti-psychotic, and anti-convulsant (Russo & Marcu, 2017). CBD also has very few demonstrated side effects, does not produce euphoria (‘high’) (Spindle et al., 2020) and has a little to no abuse liability (Babalonis et al., 2017; Viudez-Martinez et al., 2019). CBD is thus a desirable candidate molecule given its unique pharmacological profile.

In a survey of individuals undergoing in-patient treatment for OUD, nearly half (41.9%) reported using CBD to ease withdrawal symptoms (Kudrich, et al., 2024). One randomized controlled trial found that off-label oral administration of an FDA-approved pharmaceutical grade CBD formulation (Epidiolex^®^) reduced opioid craving and anxiety in humans with OUD, where effects lasted for at least a week (Hurd et al., 2019). While this adds to the promise of CBD as an OUD pharmacotherapeutic, the effects of CBD on opioid dependence and withdrawal have not yet been systematically studied (Le, Au, Hua, & Le, 2023). The limited preclinical literature demonstrates that CBD decreased naloxone-precipitated gastrointestinal and locomotor withdrawal symptoms and spontaneous withdrawal somatic signs, motor activity, and anxiety-like behaviors in opioid-dependent male and female mice (Hayduk, Hughes, Winter, Milton, & Ward, 2024; Navarrete, Gasparyan, & Manzanares, 2022; Scicluna et al., 2024). Continued research is required to evaluate and extend these findings.

This study investigated effects of CBD on acute and protracted spontaneous withdrawal symptoms in morphine-dependent male and female rats. Rats were injected twice-daily with escalating doses of morphine for 5 days and maintained at the highest does for 5 days to induce opioid dependence. Spontaneous withdrawal was then induced via abrupt discontinuation of morphine. Rats were administered CBD or its vehicle once daily via oral gavage starting 14-hr after morphine discontinuation, and withdrawal symptoms were assessed at acute (38-hr) and protracted (up to day 7) timepoints. We hypothesized that CBD would improve withdrawal symptoms, evidenced by a reduction of the acute physical symptoms (e.g., weight loss, body shakes, gastrointestinal distress), withdrawal-induced pain hypersensitivity, and protracted anxiety-like behavior.

## Methods

### Animals

Adult male and female Sprague Dawley rats (Charles River, Wilmington, MA, USA) were single housed in wire-topped, plastic cages (27 × 48 × 20 cm) with standard enrichment and *ad libidum* food and water access in the home cage. Diet was a corn-based chow (Teklad Diet 2018; Harlan, Indianapolis, IN, USA). The vivarium was on a 12-hr reverse light/dark cycle (lights off at 8:00 a.m.) and was humidity and temperature controlled. All procedures used in this study were approved by the Johns Hopkins Institutional Animal Care and Use Committee. The facilities adhered to the National Institutes of Health Guide for the Care and Use of Laboratory Animals and were AAALAC-approved.

### Drugs

Morphine sulfate was mixed in 0.09% saline vehicle and administered via subcutaneous (s.c.) injection. Hemp-derived Cannabidiol isolate was mixed in USP sesame oil as vehicle and administered via oral gavage (p.o., 1 ml/kg) at 10 mg/kg and 30 mg/kg doses. These doses were selected to approximate CBD doses available to recreational/medical users in the consumer marketplace and used for self-prescribed symptom relief (Boehnke, Gagnier, Matallana, & Williams, 2022; Kaufmann, Bozer, Jotte, & Aqua, 2023). Drugs were prepared fresh before each administration.

### Morphine administration and spontaneous opioid withdrawal

Opioid dependence in rats was induced as previously described (Houshyar, Gomez, Manalo, Bhargava, & Dallman, 2003). Rats received twice daily injections (0700-hr and 1700-hr) of escalating morphine doses (10, 20, 30, 40, 50 mg/kg/injection) administered for the first 5 days followed by maintenance of the 50 mg/kg/injection dose for 5 subsequent days (10 days total). The final morphine injection was staggered to allow each rat to undergo withdrawal testing at the same time from their last injection. After the last injection on day 10, morphine was abruptly discontinued to induce spontaneous withdrawal. At the same time each day, body weights were recorded and food in the home cage weighed (g) to determine amount eaten.

### CBD administration

CBD or vehicle was administered to rats daily, beginning 14-hr after morphine discontinuation (on ‘day 0’ of withdrawal). Assessments of withdrawal occurred 1-hr following administration of CBD or vehicle at 38-hrs (‘day 1’), after the 5^th^ treatment administration (‘day 4’), and after the 8^th^ treatment administration (‘day 7’). This pretreatment time was selected based on prior studies showing a T_max_ of 1-2 hours following oral administration of CBD (Deiana et al., 2012; Schwotzer et al., 2023). Assessments took approximately 45 minutes to complete.

### Assessment of somatic signs

Withdrawal somatic signs were evaluated at acute and protracted timepoints on days 1, 4 and 7 post-morphine discontinuation. Rats were placed in clear Plexiglas chambers (9.6 in x 9.6 in x 14.6 in) and video recorded for 10 min. Trained experimenters blinded to the experiment conditions scored videos offline using a standardized scoring protocol based on validated checklists (Gellert & Holtzman, 1978; Schulteis, Markou, Gold, Stinus, & Koob, 1994). Two experimenters scored each rat. Average inter-rater reliability scores were >90% agreement (McHugh, 2012). An average of the two scores was used as a single datapoint for each rat. Withdrawal-related behaviors were scored using Boris software (Friard & Gamba, 2016). Data were transformed to change from baseline values, and a weight was applied based on the presence/absence or frequency of the behaviors. Behaviors (with weighted score criteria) were: wet dog shakes (1-2: 2 pts; 3+: 4 pts), head shakes (1-2: 2 pts; 3+: 4 pts), escape jumps (1-4: 1 pt; 5-9: 2 pts; 10+: 3 pts), irritability on handling (3 pts.), vocalization (3 pts.), swallowing movement (2 pts), facial fasciculations (2 pts), abnormal posture (3 pts), ptosis (2 pts), chromodacryorrhea (5 pts), and diarrhea (2 pts) (Gellert & Holtzman, 1978; Schulteis, et al., 1994; Swain et al., 2020). These somatic signs encompass behaviors commonly reported in the preclinical literature that map onto human symptoms (Dunn, Huhn, Bergeria, Gipson, & Weerts, 2019; Gipson, et al., 2021).

### Von Frey test of mechanical pain sensitivity

The von Frey test was used to assess mechanical pain sensitivity during opioid dependence and withdrawal as previously described (Jenkins, Moore, & Weerts, 2025). Rats were placed in clear plastic cubicles on an elevated screen platform. Each rat hind paw was probed on the plantar surface for 3 sec with von Frey filaments (9 filaments, 0.6–15.0 g, beginning with 2.0 g). The up-down method (Chaplan, Bach, Pogrel, Chung, & Yaksh, 1994; Dixon, 1991) was used to determine the 50% paw withdrawal threshold from the individual response pattern and the force of the last von Frey filament tested. Paw withdrawal thresholds (*PWT* = 10^*xf*+*kẟ*^) were calculated for each paw and averaged to yield a single datapoint for each animal. Log-scale data from the PWT formula were computed as percent maximum possible effect (*MPE* [%] = (log (*PWT*) − log (*μ*_*sal*_))⁄(log (15) − log (*μ*_*sal*_))), where *μ*_*sal*_ is the average *PWT* for within-sex controls (Mills et al., 2012), to standardize measures to other published studies. Baseline thresholds were obtained prior to starting morphine administration. Measurements occurred on days 1 and 10 of morphine administration (30 min post-injection) and on days 1, 4, and 7 of withdrawal.

### Elevated plus maze test of anxiety-like behavior

Rats were assessed for anxiety-like behavior on day 4 of opioid withdrawal using the elevated plus maze (EPM). Rats were placed in the center of the maze facing one of two open arms and were recorded for 5 min as they explored the two open and two closed arms of the maze. Measures of arm exploration (time spent and frequency of visits) in each of the four arms were tracked using Ethovision XT 16 (Noldus Information Technology, Leesburg, Virginia, United States). Increased time spent in the open arms of the maze was used as a measure of decreased anxiety-like behavior.

### Open field test of anxiety-like behavior

Rats were assessed for anxiety-like behavior on day 7 of opioid withdrawal using the open field test (OFT) (Prut & Belzung, 2003). Rats were placed in the center of an open field arena under bright lights and measures of arena locomotion (i.e., time spent in outer vs. center zones) were tracked for 5 min using Ethovision XT 16. Increased time spent in the center of the OFT arena, or reduced thigmotaxis, was used as a measure of decreased anxiety-like behavior.

### Statistical analysis

A total of N = 100 rats (50% female) was used for this study. Sample size was determined from past studies to be powered to detect sex differences in primary outcomes (n = 8–9 per sex/group). Analyses of variance (ANOVAs) were conducted with sex (male, female), dependence group (saline, morphine), and CBD dose (0, 10, 30 mg/kg) as between-subject factors. Repeated measures ANOVAs were used to evaluate change across time (day as within-subject factor). Greenhouse-Geisser corrections were used for violations of sphericity. Post hoc testing with Holm-Bonferroni corrections were performed where main or interactive effects were significant. Data were analyzed using JASP v0.95.0. Analyses were subsequently run independently for each sex as significant effects of sex were observed with initial three-way ANOVAs. A p < 0.05 level for two-tailed tests was accepted as significant. Comparisons of primary outcomes between saline controls and morphine-dependent animals were performed within the vehicle group. Comparisons between CBD dose conditions were performed within each dependence group. Primary outcome measures were: change in body weight from start of morphine administration/withdrawal [Δ g], daily body weight change [Δ g], daily food intake [% control], and weighted composite scores of somatic signs [Σ*⍺_i_*]); pain hypersensitivity (von Frey paw withdrawal thresholds as percent maximum possible effect, MPE [%] and in grams [g]); and anxiety-like behavior (EPM open arm time [s] and visits [n]; OFT center zone time [s] and visits [n]).

### Transparency and Openness

We affirm we have reported our sample size determinations, any data exclusions (there were none), all data transformations/calculations, and all measures used in the study. We report all software packages used to analyze and visualize the data. All data, analysis code, and research materials are available upon request. This study’s design and its analysis were not pre-registered.

## Results

### Somatic signs of withdrawal during morphine administration

Opioid withdrawal behaviors (somatic signs) were scored during morphine administration for comparative purposes to characterize the onset and time course of opioid withdrawal; results indicated these scores were minimal (between 0–2; Figure S1A). A sex x morphine interaction (F[1,96] = 12.556, p < 0.001, 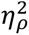 = 0.116) and main effect of day (F[1,96] = 12.095, p < 0.001, 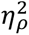 = 0.112) were observed for weighted composite (“withdrawal”) scores. In males, withdrawal scores in rats receiving morphine were lower compared to saline controls across days (main effect of morphine: F[1,48] = 4.970, p = 0.031, 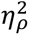 = 0.094). In females, withdrawal scores in rats receiving morphine were equal to saline controls on day 1, but increased on day 10 (compared to day 1: p < 0.001, compared to saline: p = 0.004; morphine x day interaction: F[1,48] = 6.788, p = 0.012, 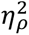 = 0.124). Overall, weighted scores on days 1 and 10 of morphine administration were all near 0 with subtle differences between rats receiving morphine and saline controls.

### Food intake and body weight during morphine administration

Food intake was suppressed during morphine administration, with more enduring effects in males (Figure S1B). A significant sex x morphine x day interaction was observed (F[7.123, 683.834] = 2.035, p = 0.047, 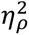 = 0.021). In males that received morphine, food intake was reduced to ∼70% of saline controls (main effect of morphine: F[1, 48] = 94.122, p < 0.001, 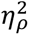 = 0.662) across all days (p’s < 0.001). In females that received morphine, food intake was initially suppressed to ∼70% of controls (morphine x day interaction: F[6.400, 307.198] = 6.170, p < 0.001, 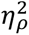 = 0.114) and steadily recovered across the first 7 days (p’s < 0.05). From day 8 onward, food intake of females that received morphine was no longer statistically lower than saline controls.

Body weight gain was also suppressed during morphine administration, more severely in males (Figure S1C). A significant sex x morphine x day interaction was observed for body weight gain during morphine administration (F[3.017, 289.619] = 40.573, p < 0.001, 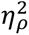 = 0.297). In males, morphine stalled weight gain (morphine x day interaction: F[2.436, 116.913] = 108.421, p < 0.001, 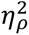 = 0.693). Significant differences from saline controls were apparent in males receiving morphine from day 4 onward (p’s < 0.05). In contrast, morphine only subtly disrupted weight gain in females. Body weights were decreased in females receiving morphine (morphine x day interaction: F[4.249, 203.939] = 13.681, p < 0.001, 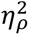 = 0.222) relative to saline controls only on day 10 (p = 0.013).

### Pain sensitivity during morphine administration

Morphine reduced mechanical pain sensitivity in both male and female rats across testing days (Figure S1D). Main effects of morphine (F[1, 95] = 201.935, p < 0.001, 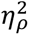 = 0.680) and day (F[1, 95] = 8.730, p = 0.004, 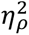 0.084) were observed. Significant sex differences were not observed. Compared to controls and baseline values, rats receiving morphine had increased paw withdrawal thresholds on days 1 and 10 of testing (morphine x day interaction for males: F[1.784, 85.632] = 35.177, p < 0.001, 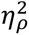 = 0.423; and females: F[1.765, 84.696] = 32.661, p < 0.001, 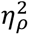 = 0.405). Thresholds were slightly decreased across timepoints in male and female saline controls (baseline vs. day 10: p’s < 0.05).

### Somatic withdrawal signs after abrupt discontinuation of morphine effects of CBD

Abrupt morphine discontinuation produced a spontaneous withdrawal syndrome in morphine-dependent male and female rats. Withdrawal symptoms (increased somatic signs and decreased food intake and body weights) were apparent one day after cessation (acute) and resolved over a 7-day (protracted) period. Daily administration of oral CBD over the 7-day period did not modify any physical symptoms of opioid withdrawal.

Morphine-dependent rats exhibited increased somatic signs in the acute phase (day 1) that subsided in the protracted phase (Figure 1A). Significant sex differences or effects of CBD were not observed. In morphine-dependent males and females, withdrawal scores were significantly increased relative to saline controls (dependence x day interaction in males: F[2,30] = 8.068, p = 0.002, 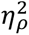 = 0.350; and females: F[2,30] = 12.883, p < 0.001, 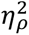 = 0.462) on day 1 for males (p < 0.001) and days 1 (p = 0.001) and 4 (p = 0.033) for females. Scores were no longer different between morphine-dependent rats and saline controls by day 7. CBD did not modify withdrawal signs in morphine-dependent males (CBD: F[2, 23] = 0.97, p=0.40, 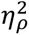 = 0.08; CBD x day: F[4, 46]=0.54, p=0.71, 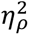 = 0.04) nor morphine-dependent females (CBD: F[2, 23] = 0.52, p=0.60, 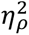 = 0.04; CBD x day: F[4, 46]=1.98, p=0.11, 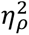 = 0.15).

**Figure 1.**
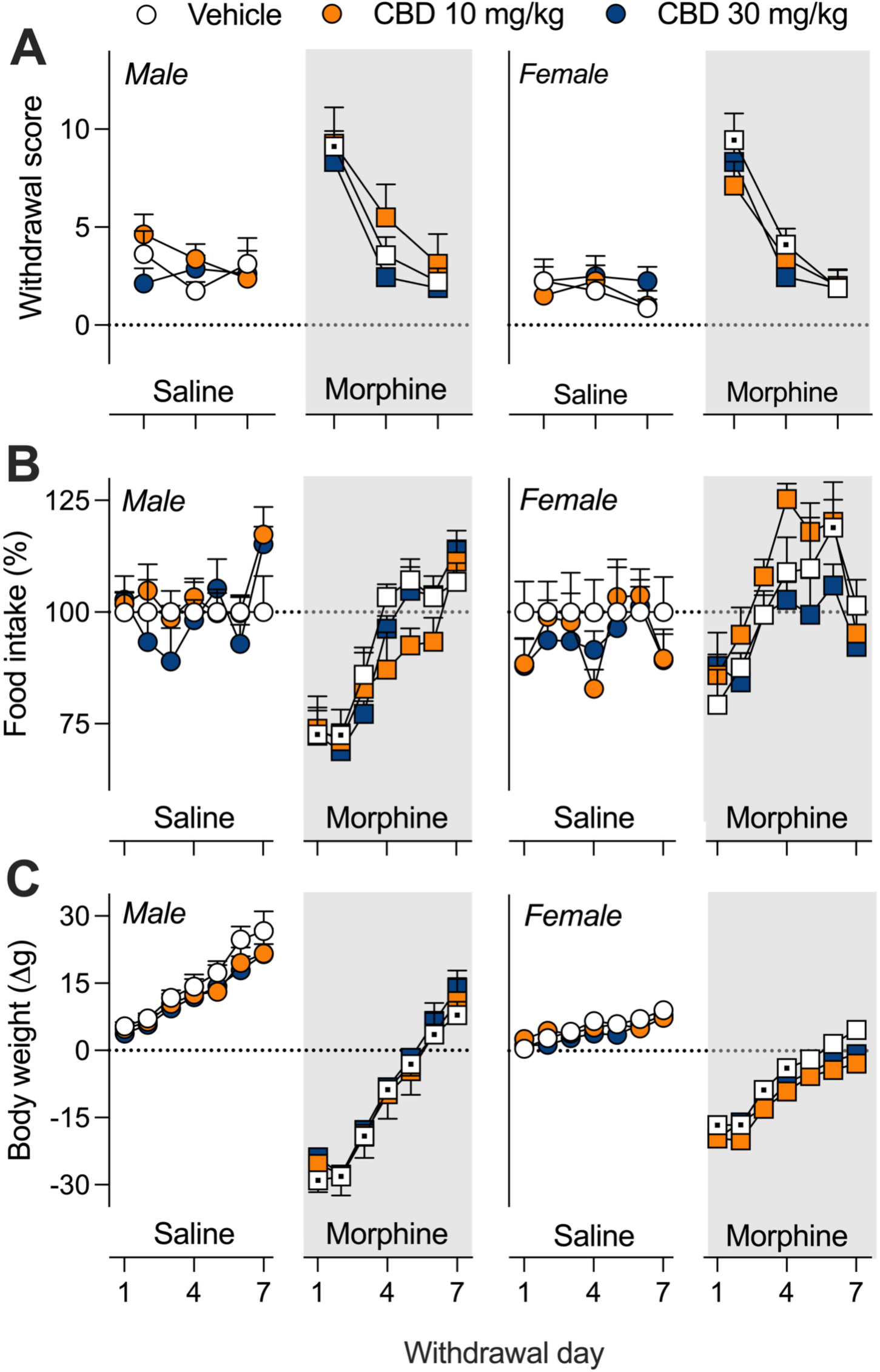
CBD effects on withdrawal score, food intake, and body weight in morphine-dependent and non-dependent rats. **A)** Daily change in body weight (Δg) from baseline (day before morphine discontinuation). **B)** Daily food intake represented as a percent of saline vehicle control values (% control). **C)** Weighted composite score of somatic signs observed on days 1, 4, and 7 of withdrawal. Data represented as means ± standard error. CBD = cannabidiol. Circle symbols = saline controls, square symbols = morphine-dependent rats. Horizontal dotted line = baseline values except for food intake and von Frey MPE where it indicates 100% and 0%, respectively. White symbols = vehicle controls, orange (light grey) symbols = 10 mg/kg CBD, blue (dark grey) symbols = 30 mg/kg CBD. White background = saline control rats, grey background = morphine-dependent rats. Symbols with center dots indicates a significant difference from saline controls (vehicle treated groups only, p < 0.05).

### Food intake and body weight loss after abrupt discontinuation of morphine

*effects of CBD* Food intake was suppressed during the acute phase of withdrawal but rebounded during the protracted phase (Figure 1B). Significant interactions of sex x dependence (F[1,96] = 2.044, p = 0.001, 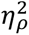 = 0.104), sex x day (F[5.345,513.110 = 11.059, p < 0.001, 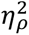 = 0.103), dependence x day (F[5.345,513.110 = 17.919, p < 0.001, 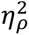 = 0.157), and sex x CBD x day (F[10.335,454.752] = 1.931, p = 0.037, 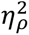= 0.042) were observed for food intake during withdrawal. Compared to saline controls, morphine-dependent males had significantly decreased food intake (dependence x day interaction: F[6,90] = 7.674, p < 0.001, 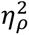 = 0.338) on days 1 and 2 (p’s < 0.05) but were no longer different than controls from day 3 onward. A significant CBD x day interaction was observed in males (F[8.711,191.633] = 2.001, p = 0.043, 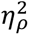 = 0.083) but differences between CBD dose conditions did not survive post hoc testing. Morphine-dependent females had no significant differences in food intake relative to controls on days 1–5 of withdrawal. Food intake was significantly increased in morphine-dependent females (dependence x day interaction: F[6,90] = 3.389, p = 0.005, 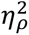 = 0.184) on day 6 (p = 0.042) before returning to control levels by day 7. Significant effects of CBD were not observed in females.

Morphine-dependent rats lost considerable body weight during the acute phase of withdrawal, which slowly recovered across the protracted phase to pre-discontinuation levels (Figure 1C). Significant interactions of sex x dependence (F[1,30] = 12.305, p = 0.001, 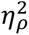 = 0.291), sex x day (F[2.701,81.017] = 14.930, p < 0.001, 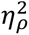 = 0.332), and dependence x day (F[2.701,81.017] = 16.505, p < 0.001, 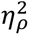 = 0.355) were observed on body weights (difference from final morphine or saline administration day; Δg). Significant effects of CBD were not observed. Relative to saline controls, morphine-dependent males and females experienced considerable weight loss (∼30g for males, ∼15g for females) at 38-hr into withdrawal (day 1; dependence x day interaction in males: F[2.184,32.758] = 9.178, p < 0.001, 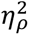 = 0.380; and females: F[3.006,45.089] = 7.611, p < 0.001, 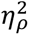 = 0.337). Body weights remained suppressed throughout withdrawal for morphine-dependent males (p’s < 0.05 on all days compared to controls) but recovered to control levels by day 5 in morphine-dependent females (days 1–4: p’s < 0.05).

### Pain sensitivity after abrupt discontinuation of morphine effects of CBD

Morphine-dependent rats showed modest increases in mechanical pain sensitivity (decreased %MPE) across the withdrawal time course (Figure 2). A main effect of dependence (F[1,88] = 13.089, p < 0.001, 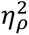 = 0.129) was observed for von Frey thresholds in an initial three-way ANOVA. Sex differences and effects of CBD were not observed. Significant differences between saline controls and morphine-dependent rats were not observed in the sex disaggregated analyses. A subsequent analysis of raw paw withdrawal thresholds also did not reveal any effects of CBD in morphine-dependent animals, however in saline controls there was a significant effect of CBD (CBD x day interaction: F[4, 84] = 2.721, p = 0.035, 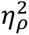 = 0.115) on raw threshold values (g). Post-hoc tests showed 10 mg/kg CBD increased thresholds on day 4 (relative to vehicle: p = 0.005; and 30 mg/kg CBD: p = 0.019) but not days 1 or 7 (Figure S2).

**Figure 2.**
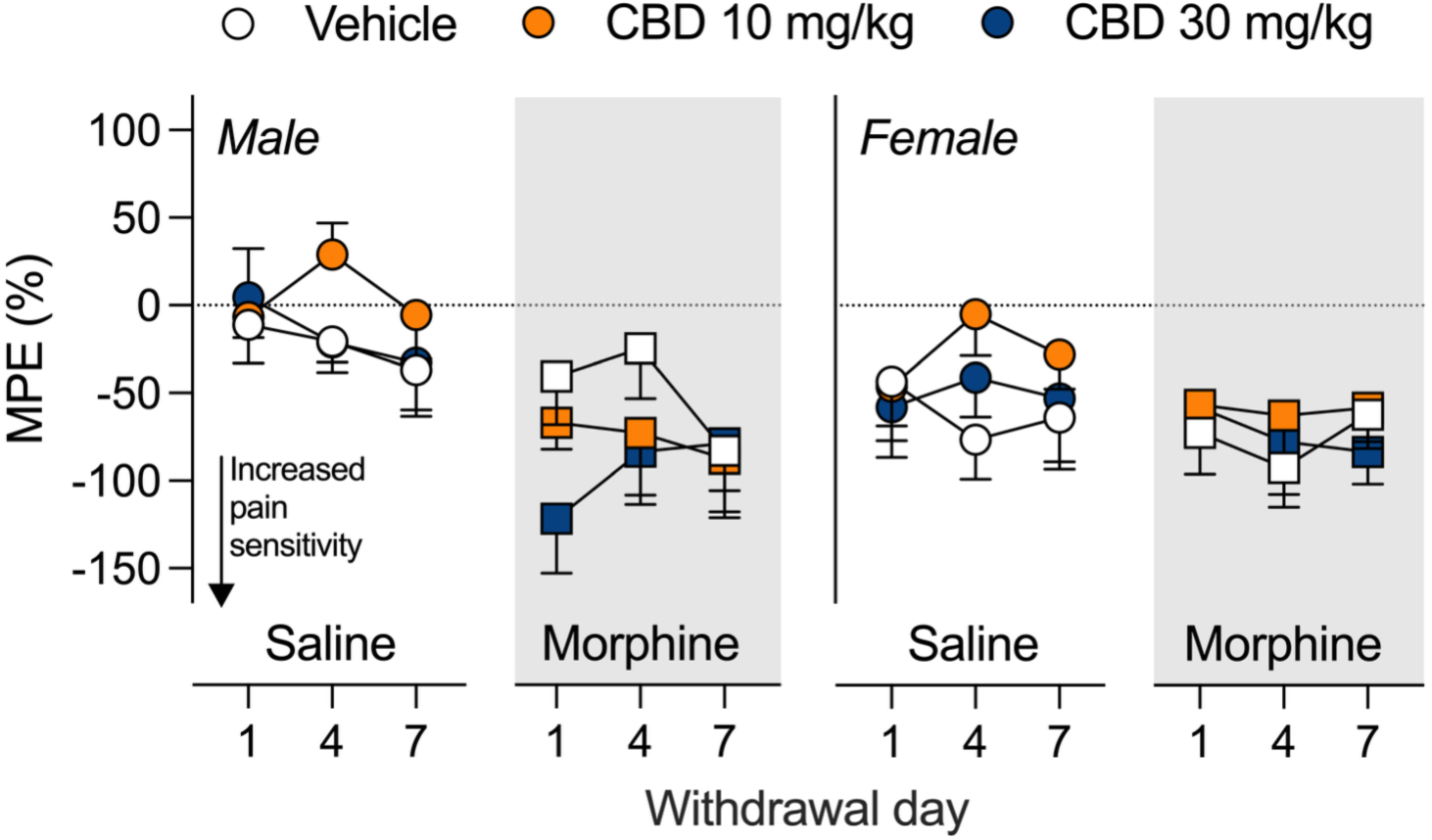
CBD effects on mechanical pain sensitivity in morphine-dependent and non-dependent rats. Von Frey paw withdrawal thresholds tested on days 1, 4, and 7 represented as a percent of the maximum possible effect (MPE, %). Data represented as means ± standard error. CBD = cannabidiol. Circle symbols = saline controls, square symbols = morphine-dependent rats. White symbols = vehicle controls, orange (light grey) symbols = 10 mg/kg CBD, blue (dark grey) symbols = 30 mg/kg CBD. White background = saline control rats, grey background = morphine-dependent rats.

### Protracted anxiety-like behavior after abrupt discontinuation of morphine effects of CBD

No differences in anxiety-like behavior were observed between saline controls and morphine-dependent rats tested during the protracted phase of spontaneous opioid withdrawal (days 4 and 7). Further, oral CBD did not affect anxiety-like behavior in either saline or morphine-dependent rats. Effects of sex were not detected in EPM primary measures on day 4 of withdrawal (Figure 3A-B). In males, differences in open arm time were not observed. Closed arm visits were decreased in morphine-dependent rats relative to saline controls (main effect of dependence: F[1, 15] = 5.467, p = 0.034, eta square = 0.267). 10 mg/kg CBD increased closed arm visits in morphine-dependent males relative to vehicle controls (p = 0,008). In females, morphine-dependent rats were not different from controls on open arm time spent or closed arm entries and CBD had no effect. On day 7 of withdrawal, effects of dependence or CBD on OFT measures were not observed (Figure 3C). A main effect of sex was observed for overall distance traveled (F[1,30] = 12.014, p = 0.002, 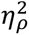 = 0.286), with females traveling greater distances than males (Figure 3D). Morphine-dependent males and females were not different from saline controls in center time or distance traveled. Significant effects of CBD were not observed in the OFT within either dependence condition.

**Figure 3.**
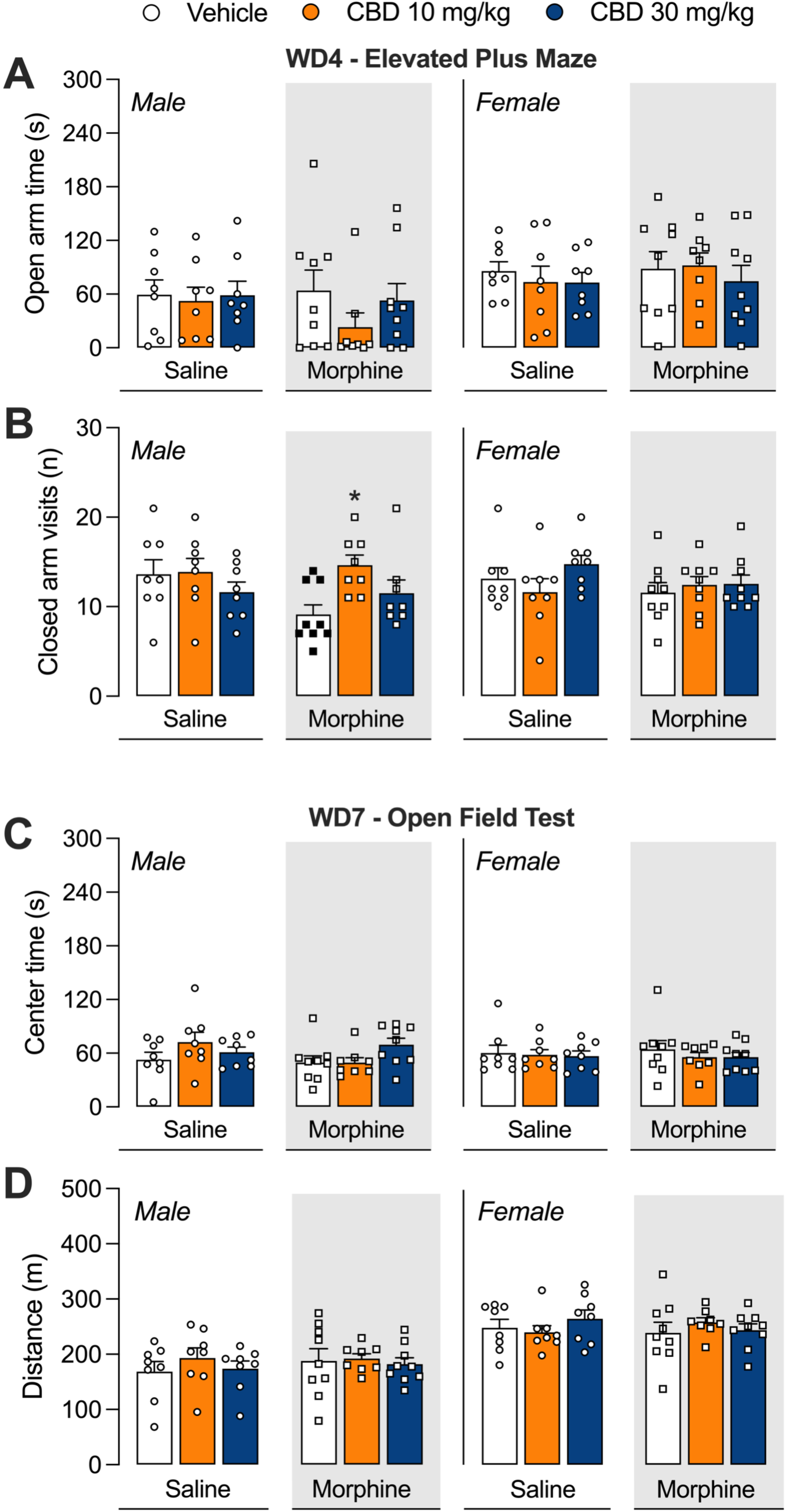
CBD effects on anxiety-like behaviors in morphine-dependent and non-dependent rats. **A)** Time spent in the open arms of the EPM. **B)** Number of visits to the open arms of the EPM. **C)** Time spent in the center zone of the OFT. **D)** Number of visits to the center zone of the OFT. Data represented as means ± standard error. EPM = elevated plus maze, OFT = open field test, CBD = cannabidiol WD = withdrawal day. Circle symbols = saline controls, square symbols = morphine-dependent rats. White bars = vehicle controls, orange (light grey) bars = 10 mg/kg CBD, blue (dark grey) bars = 30 mg/kg CBD. White background = saline control non-dependent rats, grey background = morphine-dependent rats. Closed black symbols indicates a significant difference between saline controls and morphine-dependent rats (p < 0.05).

## Discussion

This study reports that daily CBD treatment (10 or 30 mg/kg, p.o.) after abrupt discontinuation of morphine did not attenuate the acute or protracted symptoms of spontaneous opioid withdrawal in male and female rats. Following morphine discontinuation, rats demonstrated physical dependence as evidenced by decreased body weights and food intake, increased somatic signs, and pain hypersensitivity compared to saline control rats (non-dependent). Daily treatment with oral CBD in morphine-dependent rats did not prevent emergence or reduce severity of these symptoms. Anxiety-like behaviors were not detected in rats and CBD had no effect on these measures. Altogether, these data suggest that at the doses tested, oral CBD administered alone does not directly affect the physical symptoms of opioid withdrawal.

The current study was designed to model how individuals with OUD self-report using CBD to ease withdrawal symptoms or treat their anxiety and pain (Kudrich, et al., 2024), by using the most common administration route (oral) following a typical medical dosing schedule (i.e. once daily throughout withdrawal). Naturalistic and survey studies report that people using CBD for self-prescribed symptom relief take 50 mg CBD daily on average (range: 8–390 mg/day) via oral administration (Boehnke, et al., 2022; Kaufmann, et al., 2023). Doses of 10 and 30 mg/kg in a rat are roughly equivalent to 113 and 340 mg for a 70 kg human, when using a species-appropriate dose conversion (Nair & Jacob, 2016).

Recent preclinical data showed that 100 mg/kg, i.p. CBD decreased naloxone-precipitated paw tremors (Scicluna, et al., 2024) and jumping behavior (Hayduk, et al., 2024) in opioid dependent mice but not at lower doses (10, 30 mg/kg, i.p.). 10 and 30 mg/kg CBD previously decreased opioid withdrawal-related motor impairments and gastrointestinal issues in male mice (Navarrete, et al., 2022; Scicluna, et al., 2024), an effect not seen in the present study.

Differences in study outcomes may be due to differences in model species and/or administration route (oral vs i.p.) as oral doses have lower bioavailability. The present study found no beneficial effects of CBD on spontaneous withdrawal. We did find that CBD decreased pain sensitivity in saline controls on day 4 of withdrawal testing. This study adds to the existing literature reporting no effect of CBD on precipitated withdrawal symptoms, regardless of model species (Chesher & Jackson, 1985; B. Hine, M. Torrelio, & S. Gershon, 1975; Bromfield Hine, Marina Torrelio, & Samuel Gershon, 1975; Lichtman, Sheikh, Loh, & Martin, 2001).

These data do not rule out potential therapeutic effects of CBD on OUD that instead may be related to other aspects of opioid abstinence, such as cue-induced craving or negative affect. Previous preclinical data showed that CBD (5–20 mg/kg, i.p.) decreased conditioned reinstatement of heroin-seeking behavior in male rats (Ren, Whittard, Higuera-Matas, Morris, & Hurd, 2009). Epidiolex (600 mg) also decreased cue-induced craving without affecting withdrawal symptoms in patients on medication-assisted treatment, in a randomized cross-over trial (Suzuki et al., 2023) and an open-label study (Suzuki, Martin, Prostko, Chai, & Weiss, 2022). An RCT of Epidiolex (400, 800 mg) showed decreased cue-induced craving and anxiety in heroin abstinent individuals (Hurd, et al., 2019). However, positive blood screens for THC were reported in some patients, as was previously reported with Epidiolex (Johnson, Kilgore, & Babalonis, 2022; Sholler et al., 2022), making overall interpretations difficult (Hurd, et al., 2019).

Individuals with OUD will self-report using CBD and/or cannabis to ease withdrawal symptoms (Kudrich, et al., 2024), including anxiety, tremors, sleep disturbances, and gastrointestinal issues (C. L. Bergeria, Huhn, & Dunn, 2020; Meacham, Nobles, Tompkins, & Thrul, 2022).

These reports are strongly associated with respondents’ positivity toward the potential therapeutic use of CBD (Kudrich, et al., 2024). Many CBD products also contain some THC, which may contribute to effects. Thus, it is difficult to disentangle the role of THC or placebo and expectancy effects from therapeutic benefit in human studies of potential cannabinoid-based medicines (Gedin et al., 2022). Preclinical studies remain a crucial resource for systematically evaluating claims of potential therapeutic benefits from cannabinoids, to then inform continued human laboratory and clinical studies as well as public education.

The current preclinical study, which evaluated CBD effects on acute and protracted withdrawal symptoms using a clinically relevant administration route (oral dosing) and a comprehensive withdrawal symptom ethogram, is not without limitations. First, we did not evaluate blood or brain deposition of CBD in these animals. Previous work has shown that oral CBD (120 mg/kg) produced a *C_max_* in plasma and brain around 2.0 µg/ml and 8.6 µg/g, respectively, after a single dose in rats (Deiana, et al., 2012). We expect there was systemic accumulation of CBD with repeated dosing, based on existing reports of detectable amounts in plasma 24-hr after a single oral dose of 10 mg/kg CBD (Schwotzer, et al., 2023). Second, we only evaluated anxiety-like behavior in the protracted phase of withdrawal as unconditioned tests of anxiety are not repeatable. We were interested in psychological symptoms that may persist beyond somatic symptoms of withdrawal. We did not detect effects of CBD on anxiety-like behavior, as has been previously reported at 30-hr into withdrawal (Navarrete, et al., 2022). In this model, we did not observe any signs of anxiety-like behavior during the phase tested (days 4–7) but cannot rule out such effects during acute withdrawal. Future studies will investigate anxiety-like behaviors during alternative time points throughout withdrawal.

Early preclinical studies suggested that improvement in withdrawal symptoms after cannabis use is mainly driven by delta-9-tetrahydrocannabinol (THC), with CBD having a potential modulatory effect on THC (Chesher & Jackson, 1985; B. Hine, M. Torrelio, et al., 1975; Bromfield Hine, et al., 1975; Lichtman, et al., 2001). Thus, future investigations should include examination of effects of other cannabinoids, either alone or in combination with CBD. Another aspect worth investigating is whether effects relate to withdrawal severity, given differences between studies of CBD in spontaneous vs. precipitated withdrawal models. An early study showed that THC decreased precipitated withdrawal only in male mice with extended opioid exposure and more severe withdrawal (Hine, Friedman, Torrelio, & Gershon, 1975). Whether this is due to species differences, model design (e.g., differences in opioid tolerance or variability between groups) or a true dissociable effect of cannabinoids on withdrawal is not yet known.

The present study found CBD was ineffective at easing withdrawal symptoms. The current findings add to a growing literature questioning the therapeutic efficacy of CBD for treating opioid withdrawal. Indeed, prior reports assert that robust evidence considering CBD even as an adjunctive to standard opioid-replacement therapy is lacking (Kudrich, Hurd, Salsitz, & Wang, 2022). As noted in a recent umbrella review, the overall evidence for CBD alone to manage and treat substance use disorders, including OUD, is limited and inconclusive (Redonnet, Eren, Avenin, Melchior, & Mary-Krause, 2025). Concerningly, recent reports have highlighted that for some patients, the use of cannabis and/or CBD during opioid dependence or withdrawal can produce adverse effects, including worsened withdrawal symptoms (C. L. Bergeria, et al., 2020; Cecilia L Bergeria et al., 2022). However, additional studies are required to evaluate the potential therapeutic application of cannabinoid-based medicines for treating other aspects of opioid dependence and withdrawal that may improve treatment outcomes.

## Disclosures and Acknowledgements

Financial support for this work was provided by the Peter F McManus Charitable Trust (CFM). The NIDA Drug Supply Program provided the morphine for the study. Canopy Growth Corp. provided the CBD isolate for the current study. Funding sources had no other role other than financial support.

All authors have contributed to this work in an important way and have read and approved this manuscript.

CFM and EMW received funds from MyMD Pharmaceuticals, Inc. and MIRA Pharmaceuticals, Inc. for contract preclinical research unrelated to the current studies. EMW received support from Cultivate Biologics LLC, and Canopy Growth Corp. for clinical research projects unrelated to this paper.

## Authorship Contribution Statement

Conceptualization - CFM, EMW; Data Curation - BWJ, CP, RYK, CFM; Formal analysis - BWJ, CFM; Funding acquisition – CFM; Investigation - BWJ, CP, RYK, CFM; Methodology - CFM, EMW; Project administration - BWJ, CFM; Resources - CFM, EMW; Supervision - CFM, EMW; Visualization – BWJ; Writing - Original Draft – BWJ; Writing - Review & Editing - CFM, EMW

