## Supplemental Material for "Effects of oral cannabidiol (CBD) on spontaneous opioid withdrawal in male and female rats"

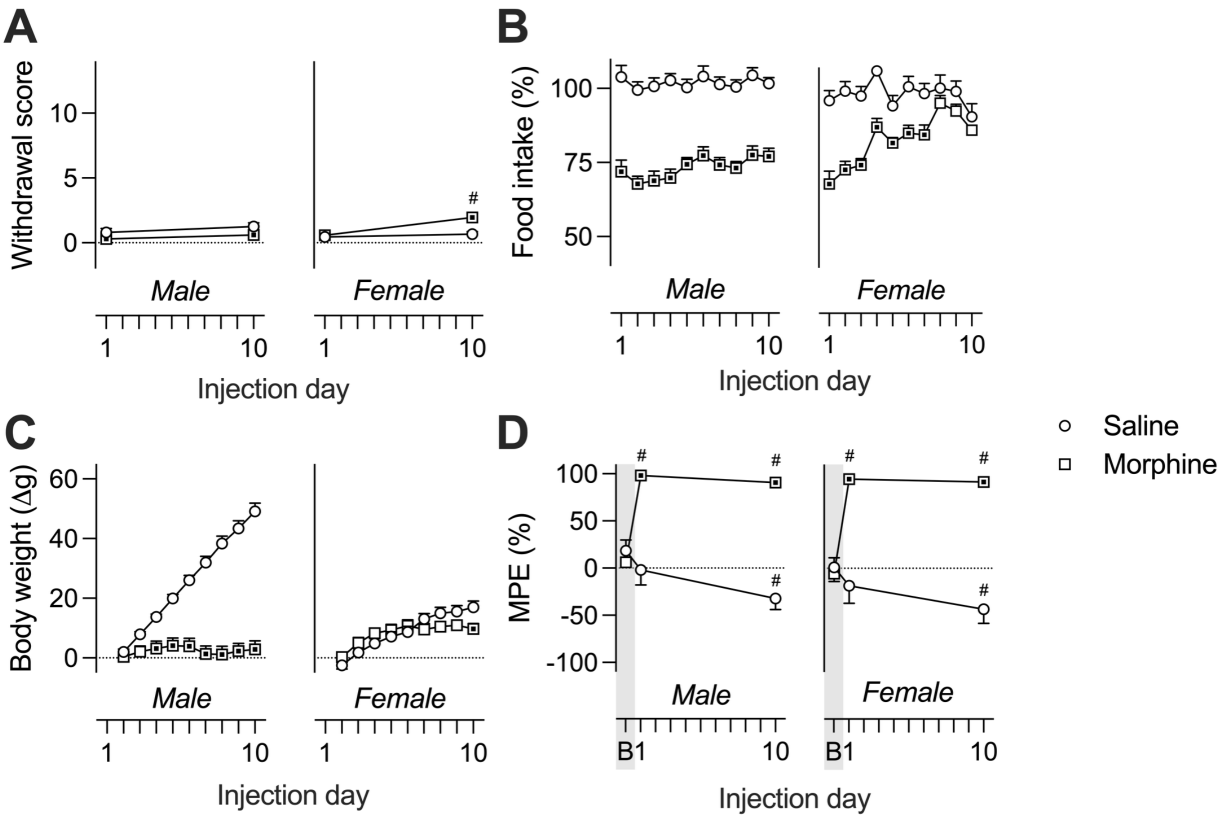


**Figure S1. Primary outcome measures during morphine administration. A)** Somatic withdrawal signs assessed on days 1 and 10 of morphine administration. **B)** Daily food intake represented as a percent of saline control values. **C)** Daily change in body weight (Δg) from baseline (before first morphine injection). **D)** von Frey paw withdrawal thresholds tested at baseline (B, grey bar) and on days 1 and 10 of morphine dependence induction, represented as a percent of the maximum possible effect (MPE, %). Data represented as means ± standard error. Circle symbols = saline controls, square symbols = morphine-dependent rats. Symbols with center dots = $p < 0.05$ between saline controls and morphine-dependent rats, × indicates a difference between males and females ($p < 0.05$). # indicates a difference from the first timepoint ($p < 0.05$).

*Change in pain sensitivity after abrupt discontinuation of morphine*

A significant CBD x day interaction was observed (F[4, 84] = 2.721, p = 0.035, $\eta_{\rho}^{2}$ = 0.115) for raw paw withdrawal thresholds within saline controls only, where 10 mg/kg CBD increased paw withdrawal thresholds on day 4 (relative to vehicle: p = 0.005; and 30 mg/kg CBD: p = 0.019) but not days 1 or 7 (Figure S2). A main effect of sex was also observed (F[1, 42] = 11.424, p = 0.002, $\eta_{\rho}^{2}$ = 0.214) where males had greater paw withdrawal thresholds than females.

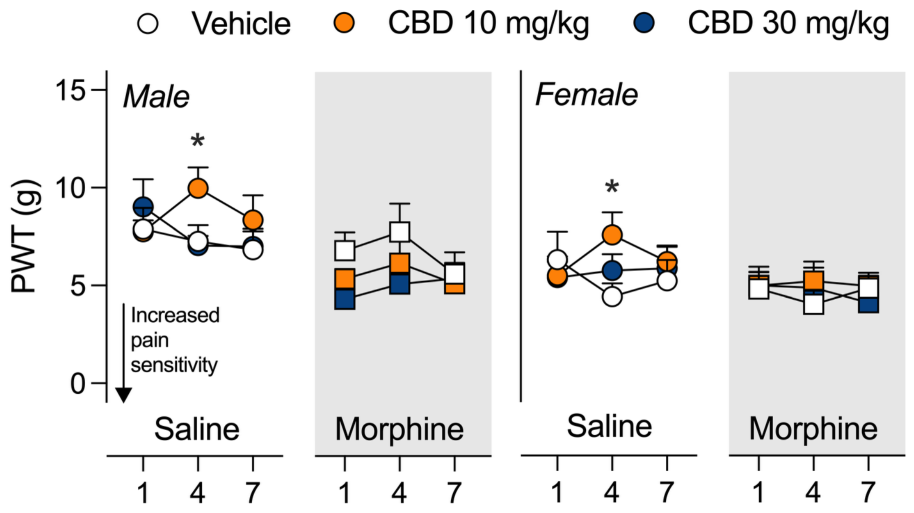


**Figure S2. von Frey paw withdrawal thresholds during opioid withdrawal.** Raw paw withdrawal thresholds measured during von Frey testing on days 1, 4, and 7 of withdrawal. Data represented as means ± standard error. CBD = cannabidiol. Circle symbols = saline controls, square symbols = morphine-dependent rats. White symbols = vehicle controls, orange (light grey) symbols = 10 mg/kg CBD, blue (dark grey) symbols = 30 mg/kg CBD. White background = saline control rats, grey background = morphine-dependent rats. * indicates difference between 10 mg/kg CBD and vehicle groups (p < 0.05).
